# Functional Quantitative Trait Loci (QTL) analysis for adaptive traits in a three-generation Scots pine pedigree

**DOI:** 10.1101/297986

**Authors:** A Calleja-Rodriguez, Z Li, H R Hallingbäck, M J Sillanpää, X Wu H, S Abrahamsson, MR García-Gil

**Affiliations:** Department of Forest Genetics and Plant Physiology, Swedish University of Agricultural Science, SE-901 83 Umeå, Sweden; Skogforsk, Box 3, SE-91821 Sävar, Sweden; Ecological Genetics Research Unit, Department of Biological Sciences, University of Helsinki, FI-00014 Helsinki, Finland.; Department of Plant Biology, Uppsala BioCenter, Linnean Centre for Plant Biology, Swedish University of Agricultural Science, SE-75007 Uppsala, Sweden; Department of Mathematical Sciences and Biocenter Oulu, University of Oulu, FI-90014, Oulu, Finland.

**Keywords:** Scots pine, climate change, quantitative trait locus, LASSO, adaptive traits

## Abstract

In forest tree breeding, QTL identification aims to accelerate the breeding cycle and increase the genetic gain of traits with economical and ecological value. In our study, both phenotypic data and predicted breeding values were used in the identification QTL linked to the adaptive value in a three-generation pedigree population, for the first time in a conifer species (*Pinus sylvestris* L.). A total of 11 470 open pollinated F_2_-progeny trees established at three different locations, were measured for growth and adaptive traits. Breeding values were predicted for their 360 mothers, originating from a single cross of two parents. A multilevel LASSO association analysis was conducted to detect QTL using genotypes of the mothers with the corresponding phenotypes and estimated breeding values (EBVs). Different levels of genotype-by-environment (G×E) effects among sites and ages were detected for survival and height. Moderate-to-low narrow sense heritabilities and EBVs accuracies were found for all traits and all sites. We identified 18 AFLPs and 12 SNPs to be associated with QTL for one or more traits. 62 QTL were significant with percentages of variance explained ranging from 1.7 to 18.9%, mostly for traits based on phenotypic data. Two SNP-QTL showed pleiotropic effects for traits related with survival, seed and flower production. Furthermore, we detected several QTL with significant effects across multiple ages, which could be considered as strong candidate loci for early selection. The lack of reproducibility of some QTL detected across sites may be due to environmental heterogeneity and QTL-by-environment effects.

## Introduction

Trees are of great economic importance as there are a source of wood and fibers in industrial processes and of immense ecologic value as carbon sinks and habitats. In the last several decades, scientists and breeders have joined efforts to improve tree health and productivity in the context of climate change, which is increasingly impacting on the adaptation of local populations.

As a result of an increase in the average temperatures in Nordic countries, longer growth seasons have been predicted which may influence adaptive traits such as timing of growth, flowering and cold acclimation of forest trees (Linderholm, 2006; Walther and Linderholm, 2006; Khorsand Rosa et al. 2015). Given the predictions of climatic change and the effects on forest fitness, tree breeders are developing strategies to breed for trees adapted to the future climatic conditions. One breeding strategy is to translocate trees (gene donors) within the species distribution range in order to introgress new gene variants (alleles) into the existing population with the aim of accelerating population adaptation (assisted gene flow) (Aitken and Whitlock, 2013). Alternatively, the populations’ standing genetic diversity could be sufficient to shift the population fitness mean towards a new optimal by artificial selection of local progenitors bearing alleles favourable to the new climatic conditions (i.e., within-population assisted selection) (Matuszewski et al. 2015). Either way, the breeding protocols involve assessment of tree performance across multiple locations as a method to evaluate and minimize genotype by environment interactions (GXE) and, thus, improve tree adaptation to an entire breeding zone (Ivkovic et al. 2015).

There are two major uses of mapping <u>Q</u>uantitative <u>T</u>rait <u>L</u>oci (QTL). The first is to accelerate the breeding cycle by artificial selection of desired genotypes at an earlier stage using molecular marker data: Marker Assisted Selection (MAS) originally introduced by Sax (1923). The breeding cycle could be shortened by the indirect selection of a causal gene (i.e., gene controlling a trait) based on the selection of molecular marker(s) in linkage disequilibrium with this gene (Collard and Mackill, 2008). The second use of genetic architecture is the development of optimal breeding methods using species genomic models (Hallingbäck et al. 2014; Wu et al. 2016).

Traditionally, QTL studies in tree species have been based on phenotypic rather than estimated genotypic or breeding values (EBVs). The main limitation of the traditional phenotypic-based QTL detection approach is the design of the experiments that typically involve only one generation, often consisting of a single full-sib family where environmental and genetic factors are confounded (Thavamanikumar et al. 2013, Isik 2014, Hall et al. 2016). Alternatively, phenotypic values can be substituted by EBVs. This method requires phenotyping progenies of the target (mother) trees and posteriorly computing EBVs with the purpose of ranking the target trees for the traits under study. Estimation of genetic effects, such as EBVs (i.e additive genetic effects), can improve the estimation of genotypic performance and also improve the genotypic value estimation of candidate genotypes in breeding programs (Piepho et al. 2008).

Since the milestone article by Lander and Botstein (1989) there have been a large number of QTL mapping studies in multiple plant species (Mauricio, 2001). In forest species, QTL analyses have either been based on progenies (full-sib or half-sib families) or unrelated individuals (association mapping). Those studies resulted in the identification of several QTL of small to moderate individual effects mostly related to growth (e.g., Lerceteau et al. 2000, 2001; Markussen et al. 2003; Yazdani et al. 2003; Bartholome et al. 2013, 2016), wood quality (e.g., Li et al. 2014; Thumma et al. 2010, Freeman et al. 2013) and disease resistance (Hanley et al. 2011). However, less effort has been devoted to the study of traits of adaptive values (Jermstad et al. 2001a,b; Hurme et al. 2000; Yazdani et al. 2003).

In this study, the goal was to identify QTL of economical and ecological value such as growth, tree quality, frost hardiness and survival measured across three different trials and multiple years in a three generation pedigree in Scots pine (*Pinus sylvestris* L.). The marker information has already been published by Li et al. (2014) but in our current study we used new growth and adaptive phenotypic traits data and EBVs to perform a novel QTL study.

## Material and Method

### Plant material

The parents (AC3065 and Y3088) of the full-sib cross are plus-trees from north Sweden (latitudes 65°08´N and 64°09´N, respectively), and belong to the Swedish breeding program (see supplementary figure S4). Progeny generated from this cross (F_1_-generation) were planted in 1988 as one year old seedlings (Abrahamsson et al. 2012). 455 fullsib individuals from this cross were included in the study (trial F485: Flurkmark, see Table 1 and supplementary figure S4). In 2006, open pollinated (OP) seeds (F_2_-generation) from 360 fullsib trees in the F485 trial were collected and grown at the Skogforsk nursery in Sävar. These fertile 360 trees producing female cones in 2006 were the only ones included in the collection as the others were non-fertile at this time. None of the 455 trees produced any male cones. The resulting OP seedlings were planted in four different field trials in 2008, using a complete randomized single-tree plot design. Three out of the four trials were chosen for this study, with an average number of trees per OP family of 9, 10 and 11 for sites F725, F723 and F726 (Table 1 and supplementary material S4), respectively (the fourth trail was unusable due to the construction of a new road over the trial). These trials could, from the technical point of view, be treated as half-sib trials. Although it should be cautioned that all F_2_-progeny were interrelated to a limited extent (single-cousins at the least).

**Table 1.**
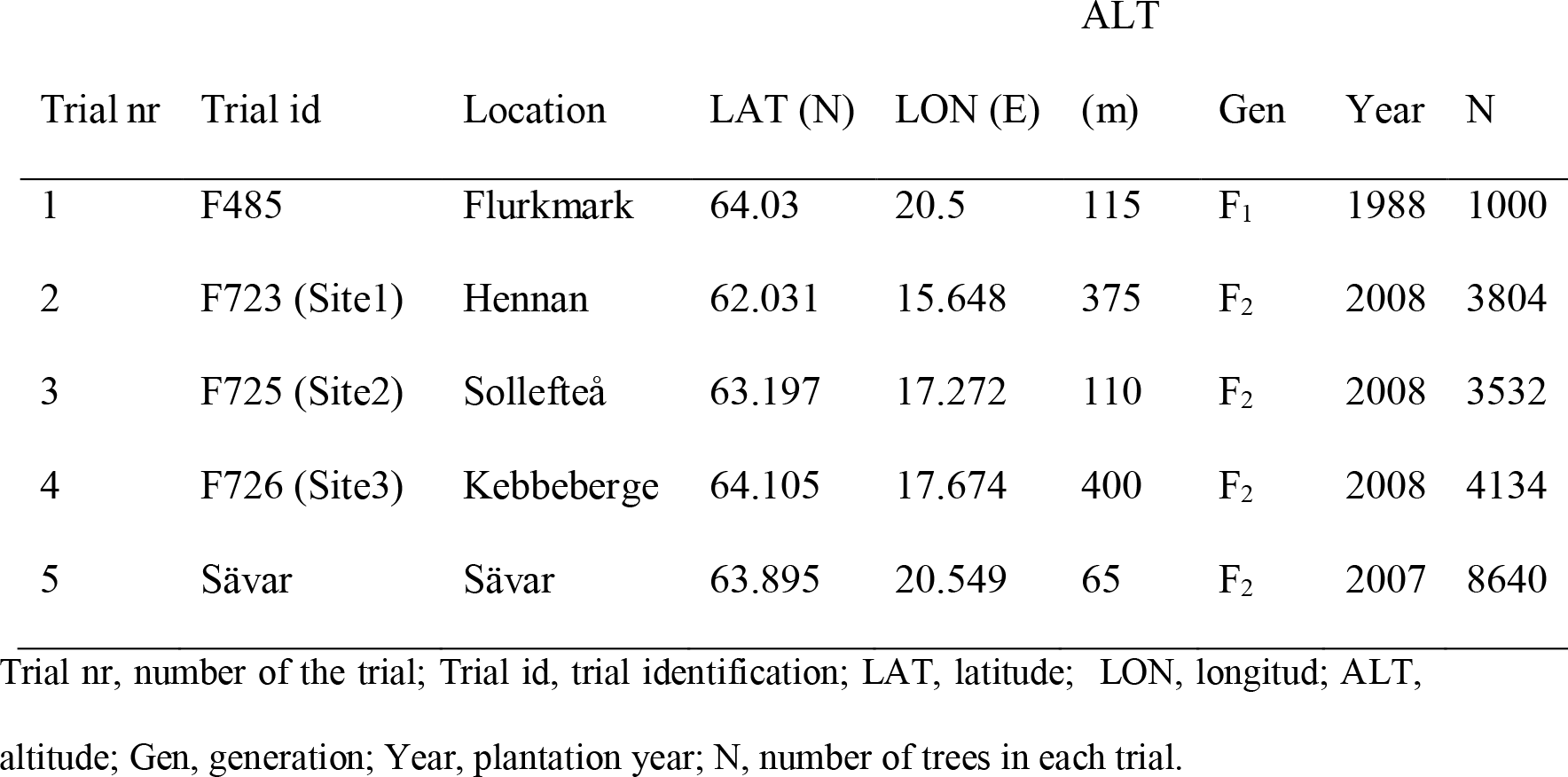
Trials description.

### Phenotypic measurements

Measured traits and the age of the trees at time of measurement for each trial are described in Table 2.

**Table 2.** Traits measured and their acronyms by trial.

| Trait | Acronym | Site |  |  |  |
| --- | --- | --- | --- | --- | --- |
|  |  | F485 | F723<br>(Site1) | F725(Site2) | F726(Site3) |
| Flower production | FP | FP |  |  |  |
| Number of cones per tree | CO | CO |  |  |  |
| Weight of 1000 seeds | W | W |  |  |  |
| Number of seeds per cone | nSC | nSC |  |  |  |
| Height | Ht | Ht_8, Ht_9, Ht_10, Ht_11, Ht_12, Ht_13, Ht_14, Ht_15, Ht_16, Ht_17, Ht_18 | eHt1_8 | eHt2_8 | eHt3_3, Ht3_8 |
| Stem diameter at breast height | DBH | DBH_8, DBH_11, DBH_16, DBH_17 | eDBH1_8 | eDBH2_8 | eDBH3_8 |
| Hardiness | Ct | Ct_4, Ct_9, Ct_11, Ct_16 |  |  | eCt3_3 |
| Branch diameter | BrD | BrD_8, BrD_11, BrD_16 |  |  |  |
| Branch quality | BrQ |  | eBrQ1_8 | eBrQ2_8 | eBrQ3_8 |
| Branch angle | BrA | BrA_8, BrA_11, BrA_16 | eBrA1_8 | eBrA2_8 | eBrA3_8 |
| Survival ability | Vt |  | eVt1_1,<br>eVt1_8 | eVt2_1, eVt2_8 | eVt3_1, eVt3_3,<br>eVt3_8 |

The first column in the table is the trait description, followed by the trait acronym used for each trait and site of measurement. The number before the underscore indicates the site where the trait was measured (except for the FS and Sävar sites where no number was used), and the number after the score indicates the age of measurement. An “e” preceding the name acronym indicates that the trait was used to calculate EBV.

#### Trait measurements on the F_1_ fullsib cross (F485)

Height (Ht) was measured annually between 1996 and 2006 using a telescopic pole from ground to terminal bud. Stem diameter at breast height (DBH) was estimated as the mean of DBH measured in two directions (north-south and east-west direction) in 1996, 1999, 2004 and 2005. Branch diameter (BrD) was scored as the horizontal diameter in two representative branches in the 10^th^ whorl counted from the top. If this branch whorl did not have any living branches, branches were chosen from the 9^th^ whorl or the next consequtive whorl with living branches. The measurements were taken at a maximum of two cm from the stem with a digital calliper, and were done in 1996, 1999 and 2004. Branch angle (BrA) was measured, in two representative branches, between the stem and the direction of the branch at the base on a scale of 360°. Branches with a representative angle of all the branches present in the tree were chosen for BrA. The two representative branches were not necessarily the same branches that were measured for BrD, but were located in the same whorl. These measurements were scored in 1996, 1999 and 2004. Ht measured between years 1996 and 1999, DBH, BrA and BrD measured at years 1996 and 1999, for individuals 1-93 have been previously published by Lerceteau et al. (2000) and Lerceteau et al. (2001).

Needles, from the trees, as close to the top as possible were collected and frozen to calculate hardiness (Ct) according to Nilsson and Walfridsson (1995) in 1992, 1997, 1999 and 2004. Presence or absence of female flower production (FP) was scored as a binary character: 1 as present and 0 as absence. Weight of 1000 seeds (W) in grams and number of cones per tree (CO) were counted exhaustively at the time of cone collection and seed extraction in 2006. The number of seeds per cone (nSC) were scored as: *nSC = seed number per tree/CO*.

#### Trait measurements on the F_2_ OP trials (F723, F725 and F726)

All phenotypic measurements are shown in Table 2 and described here. For simplicity we will refer to trial F723 as Site1, trial F725 as Site2 and trial F726 as Site3, hereafter.

Survival ability (Vt) was scored in all the three OP trials at ages one and eight plus at age three for Site3 only. Vt was scored for each tree, according to Persson and Andersson (2003), in four categorical classes: healthy, slightly damaged, severely damaged but alive, and dead. DBH, Ht and BrA were scored at age eight in all OP trials following the same protocol described for the full-sib cross (Table 2). At the same age, branch quality (BrQ) was scored in nine categorical classes considering the appearance of the entire crown in relation to the tree size and the neighbourghing trees. Finally, in Site3 Ht was measured at an additional age of three and at the same age Ct was also assessed in the manner previously described for the full-sib cross.

### DNA extraction and marker development

CTAB (Doyle, 1991) method was used to extract DNA from vegetative buds of the F_1_ full-sib individuals. All 455 individuals were genotyped by amplified length polymorphism (AFLP) markers, and a small subset of 90 individuals were genotyped for single nucleotide polymorphism (SNP) markers. Filtering and ordering resulted in a dataset of 153 AFLP markers genotyped for 455 individuals (abbreviated as A set), and a small mixed dataset of 153 AFLP and 166 SNP markers genotyped for the 90 individuals (abbreviated as S+A set). These two datasets were separately used for QTL analysis, for further details see Li et al. (2014)

### Statistical analysis for estimating breeding values (EBVs)

Progeny tests are primarily established to rank the parents according to the EBVs (White and Hodge, 2013). The different EBVs and their acronyms can be found in Table 2. An “e” preceding the trait acronym indicates that we are referring to an EBV and a number before the underscore sign refers to the OP trial. For example, eVt3_8 refers to an EBV for survival (eVt) at Site3 (eVt3) measured when the trees were eight years old (eVt3_8).

#### Single site spatial analysis

Spatial analysis is useful to detect spatial environmental variation thereby improving the quality of the adjusted data and the accuracy of the genetic parameter estimates (Cappa et al. 2015). Therefore prior to any genetic analysis the data was adjusted for within-trial environmental effects, using a two-dimensional separable autoregressive (AR1) model where row and column directions were fitted for each trial (supplementary information S2). This single model was applied to each trait data (DBH, Ht, BrQ, BrA, Vt and Ct) using ASReml3 (Gilmour et al. 2009). This method divides the residual variance into an independent component and a two dimensional spatially autocorrelated component (Dutkowski et al. 2002, 2006; Bush et al. 2013). An additional random term with one level for each experimental unit was therefore fitted ((Dutkowski et al. 2002; Ivkovic et al. 2015).

Diagnostic tools, variogram and plots of spatial residuals were used to detect design, treatment, local and extraneous effects with ASReml 3.0 (Gilmour et al. 2009). The predicted design effects and spatial residuals were extracted from ASReml output files and used to remove all environmental effects from the raw data. The adjusted data were first analysed in a single-site (univariate) analysis to estimate the genetic variance components.

#### Multi-environment analysis (MET): G×E

The spatially adjusted data were also used in a multienvironment analysis (MET). In MET the use of sophisticated models is necessary because the diverse environments may exhibit variance heterogeneity as well as genotype-by-environment interactions. These effects may affect genetic parameter estimations (Ogut et al. 2014, Isik et al. 2017). Therefore, MET was performed using a factor analytic (FA) model in order to explore the additive G×E in the best possible manner with the final objective of predicting EBVs.

FA models have become more popular and efficient in plant breeding MET (Kelly et al. 2009, Cullis et al. 2014, Smith et al. 2015, Chen et al. 2017) because they provide a good parsimonious approximation to unstructured (US) models using few informative factors.

The following individual tree (or animal) linear mixed model with FA variance structure was thus used:

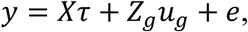

where *y* is the vector of observations for *s* sites (in our study site and trial are the same) and *m* genotypes, combined across all trials, τ is the vector of fixed effects (intercept and site) with the associated design matrix *X*; *u*_*g*_ is the random vector of genotype within environment effects (*ms* × 1) with the associated design matrix *Z*_*g*_ and *e* is the combined vector of random residuals from all sites. The random effects are assumed to follow a multivariate normal distribution with means and variances defined by: 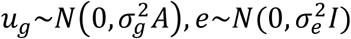

where 0 is a null vector; *A* is the numerator relationship matrix that describes the additive genetic relationships among individual genotypes; *I* is the identity matrix, with order equal to the number of trees; 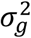 is the additive genetic variance; 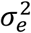 is the residual variance.

*u*_*g*_ the vector subjected to factor analysis.

FA models are usually based on the number of multiplicative terms (*k* factors) in the model (Cullis et al. 2014), so we can denote a model with *k* factors as an FAk model. Each genotype effect in each trial is a sum of *k* multiplicative terms, and we want the model that can describe most accurately the observed variance-covariance relationships among and within environments (Isik et al. 2017), using as few factors as possible in the model, in our case *k*=1. We have only 3 sites so we decided to not increase the number of *k* parameters in our study.

The mixed model in factor analytic form (see supplementary information S2) was used to model the variance structure of the additive G×E effects:

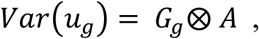

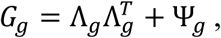

Here, *G*_*g*_ is the genetic covariance matrix among trials, *A* is the *m* × *m* numerator/additive genetic relationship matrix, Λ_*g*_ is the *s* × *k* matrix of site loadings, and, Ψ is the *s* × *s* diagonal matrix containing site-specific variance. See supplementary material for the full description of the FA model.

To perform MET, ASReml 4.0 (Gilmour et al. 2015) was used. The variance components obtained from the FA models were used to calculate narrow sense heritability and accuracy of predicted EBVs.

Narrow-sense heritability (*h*^*2*^) was calculated for each site separately but also overall across all trials, as:

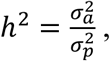

where 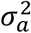 is the additive genetic variance and 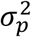 is the phenotypic variance. The accuracy of the predicted EBVs was calculated for each F_1_ full-sib mother as:

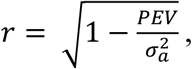

where PEV is the prediction error variance derived from the elements of the inverse of the coefficient matrix of the mixed model equations.

Further information on the models can be found in supplementary information S2.

### Statistical analysis for QTL mapping

For all the traits, each year of measurement was considered separately as a single trait and, therefore, analysed by single trait mapping approach (described below). In addition, growth trajectories were fitted by linear regression (Figure 1) to the 11 years data points (1996-2006) for Ht, and the intercept and slope parameters were taken as latent traits to be used in the subsequent QTL analysis (referred to as the two-stage approach in Li et al. 2014 to analyze longitudinal data).

**Figure 1.**
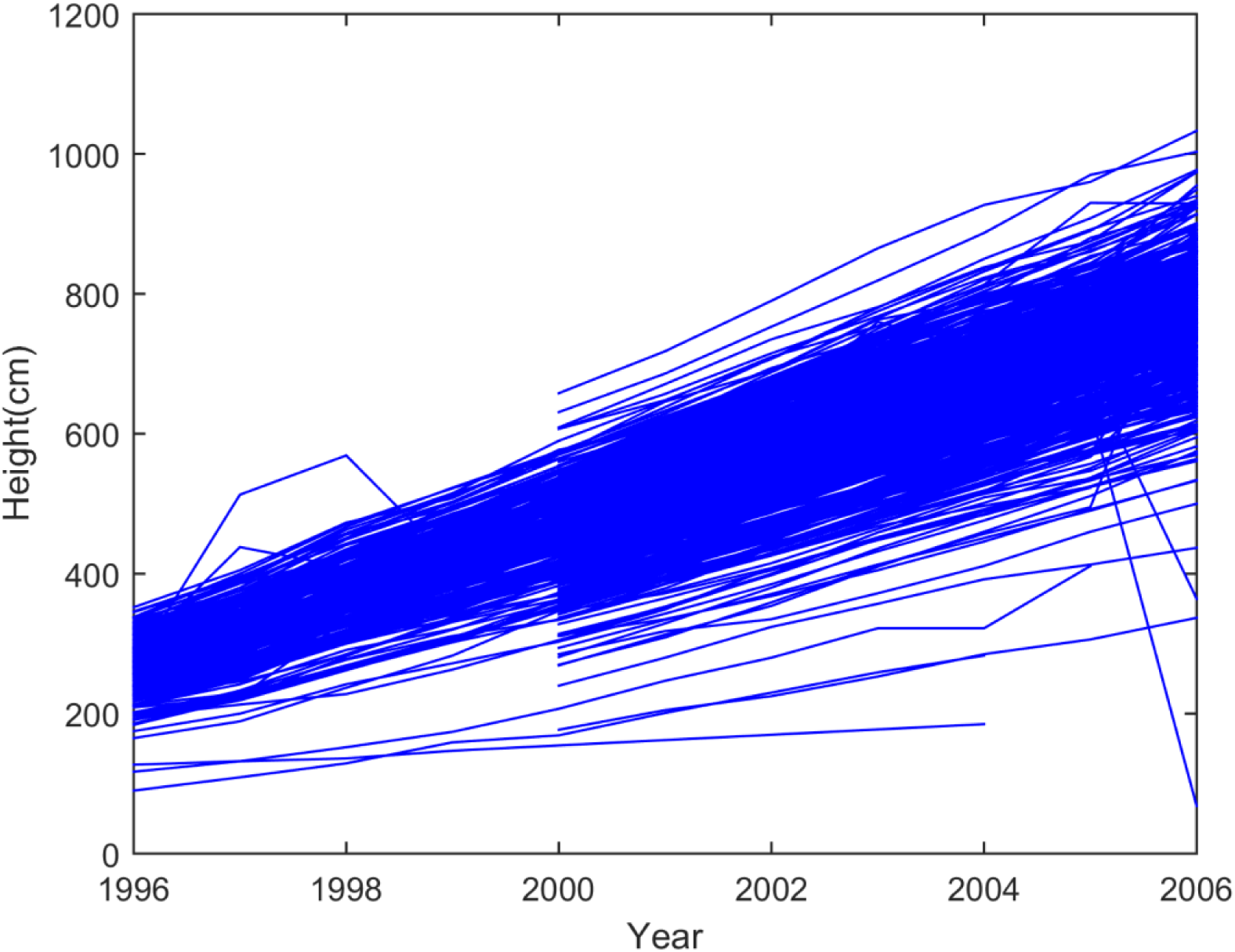
Growth trajectories of the height (measured in cm) of the 500 trees from year 1996 to 2006.

For each single trait (including each latent trait), the QTL analysis was conducted by using the LASSO regression (Tibshirani, 1996) defined by:

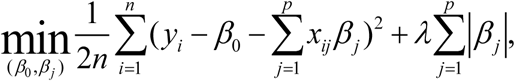

where *y*_*i*_ is the phenotypic value or estimated breeding value of individual *i* (*i*=1 … *n*; *n* is the total number of individuals), *x*_*ij*_ is the genotypic value of individual *i* and marker *j* coded as 0 and 1 for two possible marker genotypes, *β*_0_ is the population mean parameter, *β*_*j*_ is the effect of marker *j* (*j*=1 … *p*; *p* is the total number of markers). LASSO regression shrinks the coefficients of non-important markers toward zero, and only keep the important markers (i.e. those strongly associated with the traits) in the model. The tuning parameter *λ* (*λ*>0) determines how many markers should be retained in the model, and it also controls the degree of shrinkage of the regression parameters. A detailed guideline about how to choose optimal tuning parameters in the QTL mapping context can be found in Li and Sillanpää (2012). To formally judge QTL, hypothesis testing to identify QTL was conducted by a de-biased LASSO approach (Javanmard and Montanari 2014; Li et al. 2017). Due to its shrinkage nature, the original LASSO estimator do not (asymptotically) follow any parametric distribution, and therefore it is impossible to directly estimate the uncertainty of LASSO estimator such as confidence intervals and *P*-values. The De-biased LASSO approach constructed an un-biased LASSO estimator, which asymptotically follow a normal distribution. The debiased LASSO approach aims to calculate the *P*-values for all markers in the study, instead of only the markers selected by standard LASSO. Li et al. (2017) conducted a simulation study on debiased LASSO, and the method showed better power to detect QTL and stronger ability to control false positives compared to conventional single locus QTL mapping approaches. We used the following criterion for QTL judgement: a marker was claimed as a significant or suggestive QTL, if its adjusted *P*-value was smaller than 0.05 or 0.2.

## Results

### Accuracy of estimated breeding values (EBVs) and heritabilities

We calculated narrow-sense heritabilities, EBVs and accuracy of EBVs individually for each site and in the performed MET (see Table 3).

**Table 3.** (a) Average of EBV centered to the mean (a_EBV), (b) heritabilities and their standard errors in between brackets, and (c) accuracy of the EBV (r_EBV), for the traits used both in the MET analysis and single-site analysis.

| Site/Trait | eVt_1 | eVt_3 | eHt_3 | eCt_3 | eDBH_8 | eBrQ_8 | eBrA_8 | eHt_8 | eVt_8 |
| --- | --- | --- | --- | --- | --- | --- | --- | --- | --- |
| Site1 | 2.25 |  |  |  | 29.56 | 4.43 | 4.62 | 253.98 | 2.09 |
| Site2 | 2.79 |  |  |  | 33.64 | 5.27 | 5.05 | 302.64 | 2.67 |
| Site3 | 2.88 | 2.88 | 32.56 | 15.08 | 21.06 | 4.19 | 4.42 | 219.04 | 2.18 |
| MET | 2.64 |  |  |  | 28.09 | 4.63 | 4.70 | 258.58 | 2.31 |

(b) $h^2$
| Site/Trait | Vt_1 | Vt_3 | Ht_3 | Ct_3 | DBH_8 | BrQ_8 | BrA_8 | Ht_8 | Vt8 |
| --- | --- | --- | --- | --- | --- | --- | --- | --- | --- |
| Site1 | 0.15 (0.07) |  |  |  | 0.17 (0.09) | 0.09 (0.09) | 0.03 (0.09) | 0.18 (0.09) | 0.11 (0.07) |
| Site2 | 0.10 (0.07) |  |  |  | 0.35 (0.08) | 0.10 (0.08) | 0.04 (0.07) | 0.56 (0.09) | 0.12 (0.07) |
| Site3 | 0.05 (0.06) | 0.06 (0.06) | 0.21 (0.07) | 0.55 (0.09) | 0.07 (0.08) | 0.19 (0.08) | 0.22 (0.08) | 0.10 (0.08) | 0.42 (0.07) |
| MET | 0.10 |  |  |  | 0.16 | 0.06 | 0.06 | 0.24 | 0.11 |

(c) r\_EBV
| Site/Trait | eVt_1 | eVt_3 | eHt_3 | eCt_3 | eDBH_8 | eBrQ_8 | eBrA_8 | eHt_8 | eVt_8 |
| --- | --- | --- | --- | --- | --- | --- | --- | --- | --- |
| Site1 | 0.28 |  |  |  | 0.25 | 0.19 | 0.12 | 0.27 | 0.25 |
| Site2 | 0.23 |  |  |  | 0.37 | 0.22 | 0.14 | 0.44 | 0.25 |
| Site3 | 0.18 | 0.19 | 0.32 | 0.44 | 0.18 | 0.29 | 0.31 | 0.21 | 0.44 |
| MET | 0.32 |  |  |  | 0.41 | 0.29 | 0.28 | 0.47 | 0.43 |

Moderate to low heritabilities and EBV accuracies were found for all traits in all sites. Higher accuracies were found for Ht_8 (0.47) and DBH_8 (0.41) using the MET analysis (Table 3). From the single-site analysis point of view, Site2 showed the highest accuracies for DBH_8 (0.37) and Ht_8 (0.44). The EBV accuracy of Ht at Site3 decreased from 0.32 to 0.21 for age three and eight, respectively. The accuracy of survival ability at Site3, Vt3_8 (0.44), was the highest accuracy among all sites and it was almost equal to the accuracy estimated with the MET for Vt_8 (0.43). In the case of Vt_1, the highest accuracy was estimated through the MET (0.32), where single site analysis resulted in slightly smaller estimates (0.18 to 0.28). EBV accuracy of Ct at Site3 age three was moderate (0.44).

Heritabilities at Site2 (Table 3) were only higher than for the other sites for DBH_8 (0.35) and Ht_8 (0.56). Among all sites, Site3 had the highest heritabilites for BrQ3_8 (0.19), BrA3_8 (0.22), and Vt 3_8 (0.42). The heritability of hardiness measured at age three was 0.55. The heritabilities obtained for the remaining traits and sites were low, ranging between 0.03 and 0.10 (Table 3). Heritabilities estimated with the MET analysis were low for all traits, probably due to the considerable G×E-interactions detected in some cases (Table 3) and are described in the following section.

### Multi-environment analysis (MET): G×E

To detect G×E interaction patterns, genetic correlations across sites were used and are shown in Table 4. The genetic correlations for Ht_8 and DBH_8 (0.94 in both cases) between Site1 and Site2 were high, while the Ht_8 and DBH_8 genetic correlations between Site1 and the other sites were very low (0.26 to 0.32). For Vt at age one, low genetic correlations (0.24) were detected between Site1 and Site2, as well as between Site2 and Site3. In the case of Vt at age eight, a moderate correlation was detected between Site1 and Site3 (0.67), but otherwise low correlations were observed (0.15 to 0.24). A moderate BrQ_8 genetic correlation was detected between Site1 and Site3, and a low correlation was detected for the same trait between Site1 and Site2. However, in both cases the standard erros were very high, which made us consider these correlations as not significant. In the remaining traits all the genetic correlations were very low (below 0.10). BrA_8 genetic correlation between Site1 and Site2 could not be estimated.

**Table 4.** Across site genetic correlations (and standard errors) for traits used in the MET analysis.

|  | Vt_1 |  | Vt_8 |  | DBH_8 |  | Ht_8 |  | BrQ_8 |  | BrA_8 |  |
| --- | --- | --- | --- | --- | --- | --- | --- | --- | --- | --- | --- | --- |
|  | Site2 | Site3 | Site2 | Site3 | Site2 | Site3 | Site2 | Site3 | Site2 | Site3 | Site2 | Site3 |
| Site1 | 0.24±0.36 | 0.99±0.73 | 0.24±0.29 | 0.67±0.24 | 0.94±0.30 | 0.28±0.36 | 0.94±0.25 | 0.32±0.28 | 0.15±0.54 | 0.43±0.45 | n.e | 0.09±0.33 |
| Site2 |  | 0.24±0.34 |  | 0.15±0.20 |  | 0.26±0.34 |  | 0.30±0.26 |  | 0.07±0.33 |  | 0.09±0.32 |

n.e.: non estimable.

### QTL detection

We found 18 AFLPs and 12 SNPs associated across all phenotypic traits, intercepts and EBVs (Table 5 and supplementary S3). 62 QTL were significant with a range of %PVE explained from 1.7 to 18.9 (Table 5 and supplementary table 1). The lowest %PVE were detected for CO (1.70), followed by Ht at different ages (%PVE from 1.90 to 3.60), Ht intercept (2.00), FP (2.40), DBH (2.20 - 6.40), BrA (3.4), eBrA (3.4), eVt (4), eHt (4.30), Ct (4.30), DBH (5.70), Ht (5.92) and eCt (7.20).

**Table 5.**
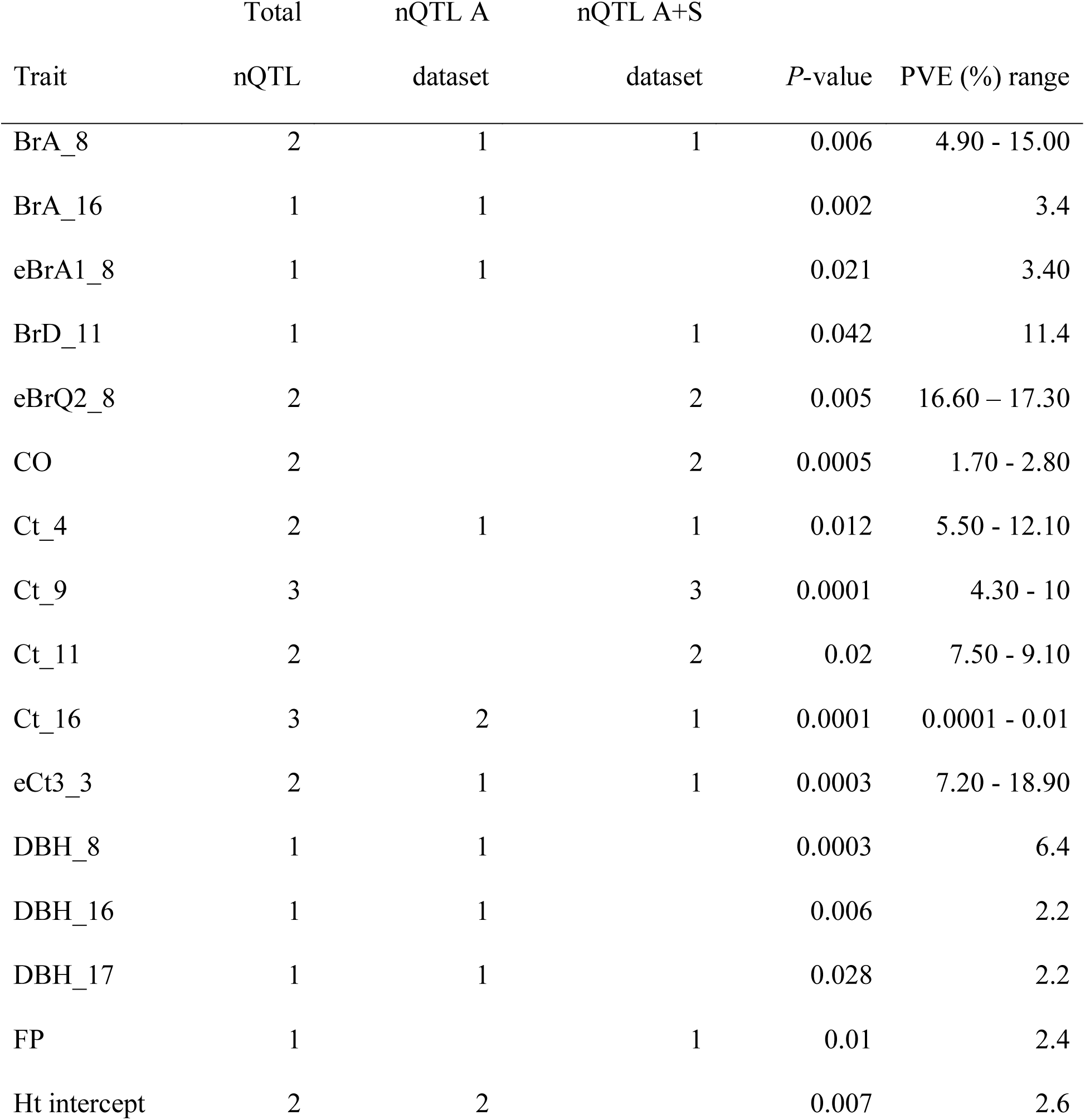
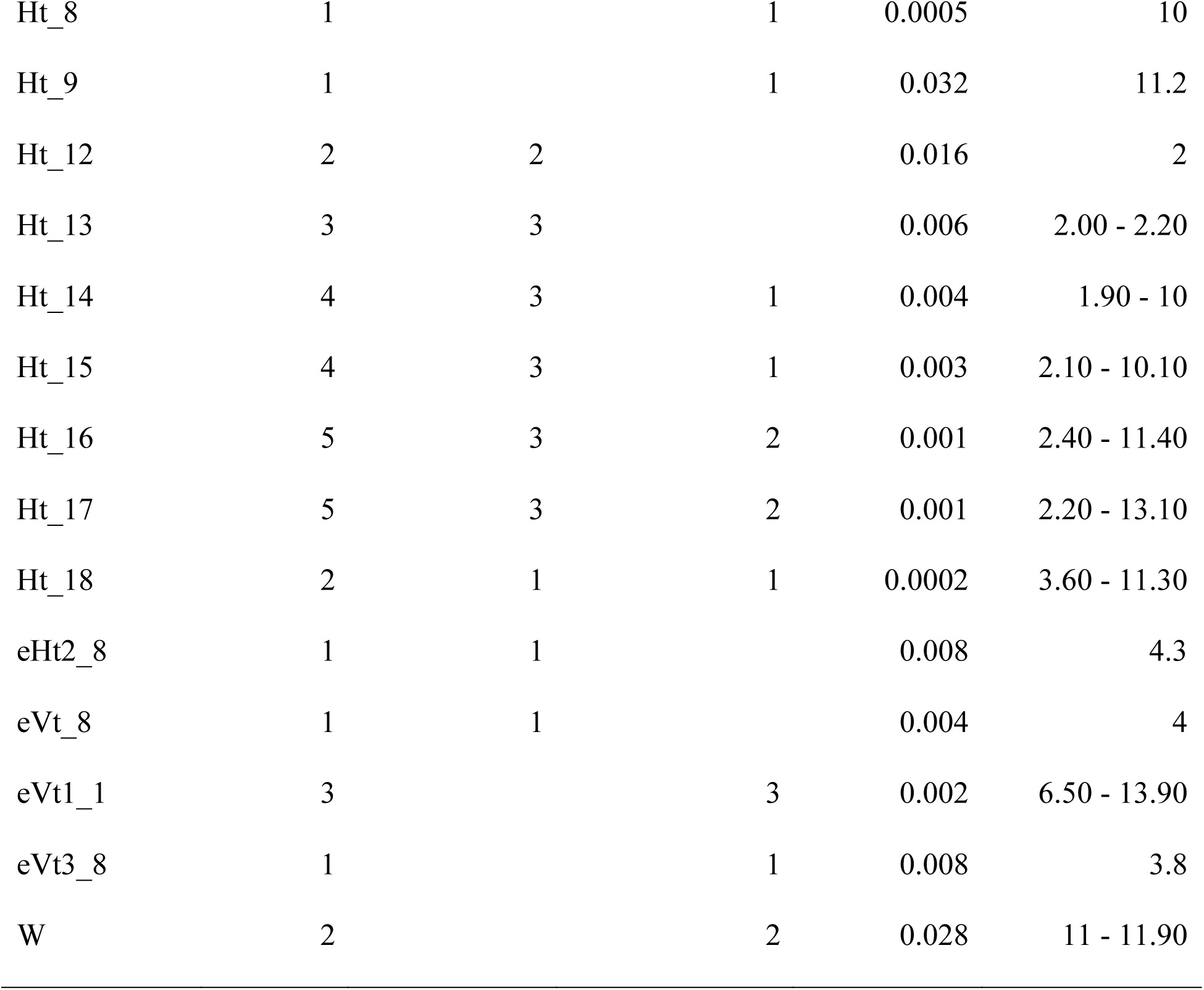
Description of the significant QTL.

| Trait | Total<br>nQTL | nQTL A<br>dataset | nQTL A+S<br>dataset | *P*-value | PVE (%) range |
| --- | --- | --- | --- | --- | --- |
| BrA_8 | 2 | 1 | 1 | 0.006 | 4.90 - 15.00 |
| BrA_16 | 1 | 1 |  | 0.002 | 3.4 |
| eBrA1_8 | 1 | 1 |  | 0.021 | 3.40 |
| BrD_11 | 1 |  | 1 | 0.042 | 11.4 |
| eBrQ2_8 | 2 |  | 2 | 0.005 | 16.60 – 17.30 |
| CO | 2 |  | 2 | 0.0005 | 1.70 - 2.80 |
| Ct_4 | 2 | 1 | 1 | 0.012 | 5.50 - 12.10 |
| Ct_9 | 3 |  | 3 | 0.0001 | 4.30 - 10 |
| Ct_11 | 2 |  | 2 | 0.02 | 7.50 - 9.10 |
| Ct_16 | 3 | 2 | 1 | 0.0001 | 0.0001 - 0.01 |
| eCt3_3 | 2 | 1 | 1 | 0.0003 | 7.20 - 18.90 |
| DBH_8 | 1 | 1 |  | 0.0003 | 6.4 |
| DBH_16 | 1 | 1 |  | 0.006 | 2.2 |
| DBH_17 | 1 | 1 |  | 0.028 | 2.2 |
| FP | 1 |  | 1 | 0.01 | 2.4 |
| Ht intercept | 2 | 2 |  | 0.007 | 2.6 |
| Ht_8 | 1 |  | 1 | 0.0005 | 10 |
| Ht_9 | 1 |  | 1 | 0.032 | 11.2 |
| Ht_12 | 2 | 2 |  | 0.016 | 2 |
| Ht_13 | 3 | 3 |  | 0.006 | 2.00 - 2.20 |
| Ht_14 | 4 | 3 | 1 | 0.004 | 1.90 - 10 |
| Ht_15 | 4 | 3 | 1 | 0.003 | 2.10 - 10.10 |
| Ht_16 | 5 | 3 | 2 | 0.001 | 2.40 - 11.40 |
| Ht_17 | 5 | 3 | 2 | 0.001 | 2.20 - 13.10 |
| Ht_18 | 2 | 1 | 1 | 0.0002 | 3.60 - 11.30 |
| eHt2_8 | 1 | 1 |  | 0.008 | 4.3 |
| eVt_8 | 1 | 1 |  | 0.004 | 4 |
| eVt1_1 | 3 |  | 3 | 0.002 | 6.50 - 13.90 |
| eVt3_8 | 1 |  | 1 | 0.008 | 3.8 |
| W | 2 |  | 2 | 0.028 | 11 - 11.90 |

Most of the significant QTL were detected for Ht phenotypic traits at different ages, followed by Ct, eVt, eCt, eBrQ, eBrA, BrA, W, CO, eHt, DBH, BrD and FP.

Eight QTL-AFLPs (TCG89, AGC91, ATG226, AGC205, TAG62, GGC205 and AGG498) were detected as singnificant, only in the A data set. The QTL-TCG89 was significantly associated to DBH at ages 16 and 17, Ht intercept and Ht between ages 12 and 18 (with *P-*values that vary from 0.011 to 0.0002).

QTL-GAT81 was significantly associated to Ht between ages 13 and 17, QTL-AGC91 to DBH at age 8. Whilst QTL-ARG226 and -AGC205 were significantly linked to BrA at ages 16 and eight respectively. QTL-TAG62 was singnificantly linked to Ct_4 with all the *P*-values ranging from 0.014 to 0.0003.

QTL-GGC205 was significantly associated to eHt2_8 (*P*-value of 0.008) and QTL-AGG498 was significant for eVt3_8 and eVt_8, with *P*-values of 0.004 and 0.008, respectively.

Five AFLP (TGG57, TCG134, TTG454, GGC97 and TTG82) were significantly associated with the S+A data sets. Those QTL were detected for BrA_8, Ct_9, eVt1_1, eVt1_1 and eBrQ2_8, respectively.

Among all the detected QTL, TCG89, GGG312, 0_7009_01-358 and axs_47-502, showed significant pleiotropic effects on DBH/Ht, eBrA1_8/W, FP/W/Ct_16/eCt3_3 and CO/Ct, respectively. AGC106, GGG312 and 0_7009_01-358 were the only three QTL significantly detected in both, the full-sib and in the half-sib progenies. AGG498 was the only QTL-AFLP showing significant association with eVt based on both, the individual and the MET, EBV estimations.

Multiple markers (CGT193, AAG115, TCG89, GAT81, AGC106, 0_7009_01-358, 2_10352_02-413, 2_9603_01-344 and 2_10212_01-241) were both significantly associated to the same trait in several years for traits such as Ht, DBH and Ct. Finally, the QTL AAG115 and TCG89 were both significantly associated with Ht and Ht_intercept, respectively.

The significant QTL detected for the EBV-based traits were only detected at one of the sites or shared between one of the sites and the MET. For the EBV of Vt, we have found four significant QTL. The QTL-AGG498 is the only case where a QTL was detected both in the single site and MET (eVt_8 at Site3 and MET). The remaining three QTL were detected only for eVt_1 at Site1 (QTL-TTG454, QTL-GGC97 and QTL-GI_F1_334). Two QTL were detected for the EBV of BrQ_8 only in Site2 (QTL-TTG82 and QTL-LP2_625).

Multiple QTL were detected for the same trait across several ages. For example, 2_10352_02-413 explains Ht variation at ages 14 to 17 (Ht_14-17). Similarly, 2_9603_01-344 is associated with Ht at ages 9 and 18, AAG115 from age 12 to 17, GAT81 from age 13 to 17 and TCG89 from age 12 to 18. Furthermore, TCG89 also explains the variation of DBH at ages 16 and 17. The SNP-QTL 0_7009_01-358 is significantly associated to Ct based on phenotypic measurements in the F_1_-generation trees at the age of 16 (Ct_16) and on EBVs scored in the F_2_ progenies at the age of 3 (eCt3_3). Similarly, AGC106 explains the observed variation for Ct at ages 9 and 16 based on phenotypic values and for age 3 based on EBVs.

## Discussion

To our knowledge, this is the first study performing an EBV-based QTL mapping in conifers. The study was also designed as a multi-environment analysis *in sensu stricto*, therefore, EBVs were computed based on pedigree adjusted values with the environmental effects having been removed.

### Genetic architecture (number and %PVE)

This QTL architechure reveals that in both data sets (A, S+A), there are more phenotypic-QTL detected than EBV-QTL. This could be explained, to some extent, by the larger number of phenotypic traits assessed compared to the EBV-based traits (30 phenotypic traits versus 21 EBV-based traits). Multiple studies on simulated and empirical data has shown that a minimum population size is required to detect QTL consistently (Hall et al. 2016), and the number of QTL detected typically increases with population size and heritability (e.g., Vales et al. 2005; Falke and Frisch, 2011; Wang et al. 2012). However, the number of QTL detected and their effects is not only a function of population size or its underlying genetic architecture (i.e., number of genes controlling the trait and their effect size). Other factors, such as marker type (dominant versus co-dominant) can also contribute to the number and effect of the detected QTL. In our study, S+A data set has similar number of AFLP and SNP markers, however, SNP-QTL represent almost the 70% (AFLP 30%) of the total number of QTL detected in the S+A data set. This is possibly due to the higher informative (codominant) nature of the SNPs (i.e., the heterotizgote Aa can be distinguished from the dominant homozygote AA).

Congruent with the theoretical and empirical expectations, our experimental design mainly allowed the detection of QTL with medium-to-large effects. In both data sets, QTL for eCt3_3 show the highest %PVE (7.20 for the A dataset and 18.90 for the S+A dataset), and exhibits the second highest *h*^*2*^ (0.55). However, in the S+A data set, we suspect %PVEs to be overestimated due to a lower number of study trees (i.e., 91 F1 trees for the phenotypic-QTL or 61 F_1_ trees for the EBV-QTL in the A+S data set compared to 496 F_1_ trees for the phenotypic-QTL or 356 F_1_ trees for the EBV-QTL in the A data set). Furthermore, Ht in Scots pine has been reported to be under low genetic control (Abrahamsson et al. 2012), in the S+A data set, the QTL associated with Ht are among the highest %PVE values.

Ct was previously known to be under moderate to high genetic control, while traits like Ht, DBH and BrA are typically described to have moderate-to-low *h*^*2*^ in multiple conifer species. For example, in *Cunninghamia lanceolata* (Lamb) Hook (Chineese fir), DBH and Ht were described to have low *h*^*2*^ (0.14 and 0.20, respectively) (Bian et al. 2014). In *Pinus elliottii* (Slash pine), low narrow-sense heritabilities were also reported for Ht (0.03), DBH (0.02) and BrA (0.06 and 0.17 depending on the population) (Pagliarini et al. 2016). Similary in *Pinus banksiana* Lamb (Jack pine) *h*^*2*^ for BrA was aslo reported to be low (0.16) (Weng et al. 2017). In *Pinus pinaster* (Ait) *h*^*2*^ for Ct estimate was higher than 0.6 (Prada et al. 2014). In *Pseudotsuga menziesi*, Ht *h*^*2*^ ranged from 0.03 to 0.09, while Ct was shown to be under low to moderate genetic control (0.16 - 0.37) (Hawkins and Stoehr, 2009). In Scots pine, Ht and Ct were reported to be under low (0.16) and moderate (0.37) genetic control, respectively (Persson et al. 2010; Abrahamsson et al. 2012).

The number of QTL and their %PVE agree with what has previously been reported in the literature (Tables 5 and 6). Several QTL have been detected for hardiness in different species such as Sitka spruce (Holliday et al. 2010), Scots pine (Hurme et al. 2000: Yazdani et al. 2003) and Coastal Douglas-fir (Jermstad et al. 2001b; Wheeler et al. 2005), with %PVE that varies from 0.7 to 24.9 depending on the specie, age and tests peformed. %PVE ranged from 1.3 to 34.9 for height at different ages in different species, for instance in white spruce (Pelgas et al. 2011), *Quercus robur* (Gailing et al. 2008; Scotti-Saintagne et al. 2004), *Eucalyptus urophylla* (Gion et al. 2011) and Scots pine (Lerceteau et al. 2000, 2001). In the case of BrD and BrA there are not many QTL studies available, but Lerceteau et al. (2001) found two and one QTL with %PVE estimates of 29.5 and 16.1 respectively.

**Table 6.**
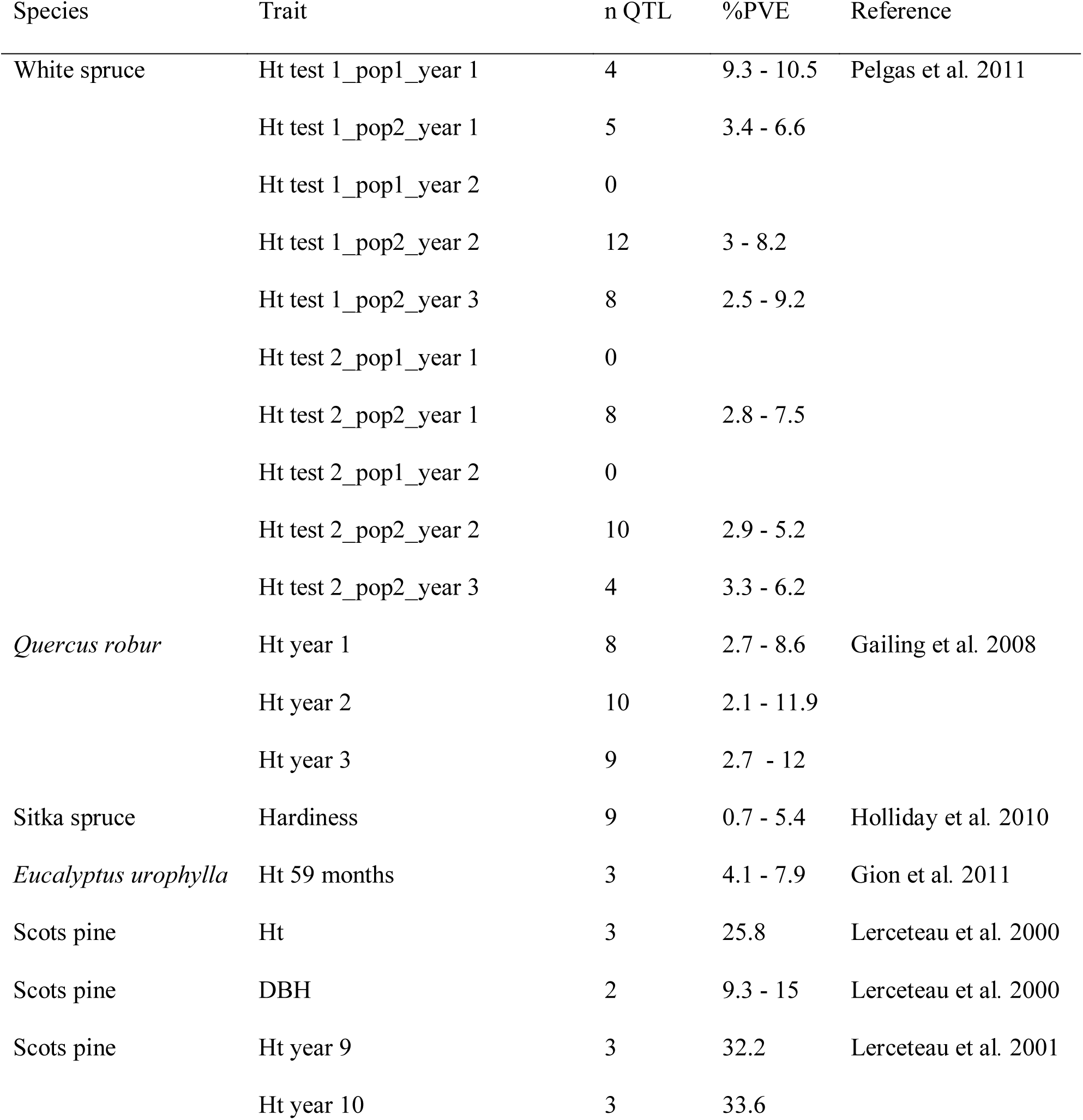
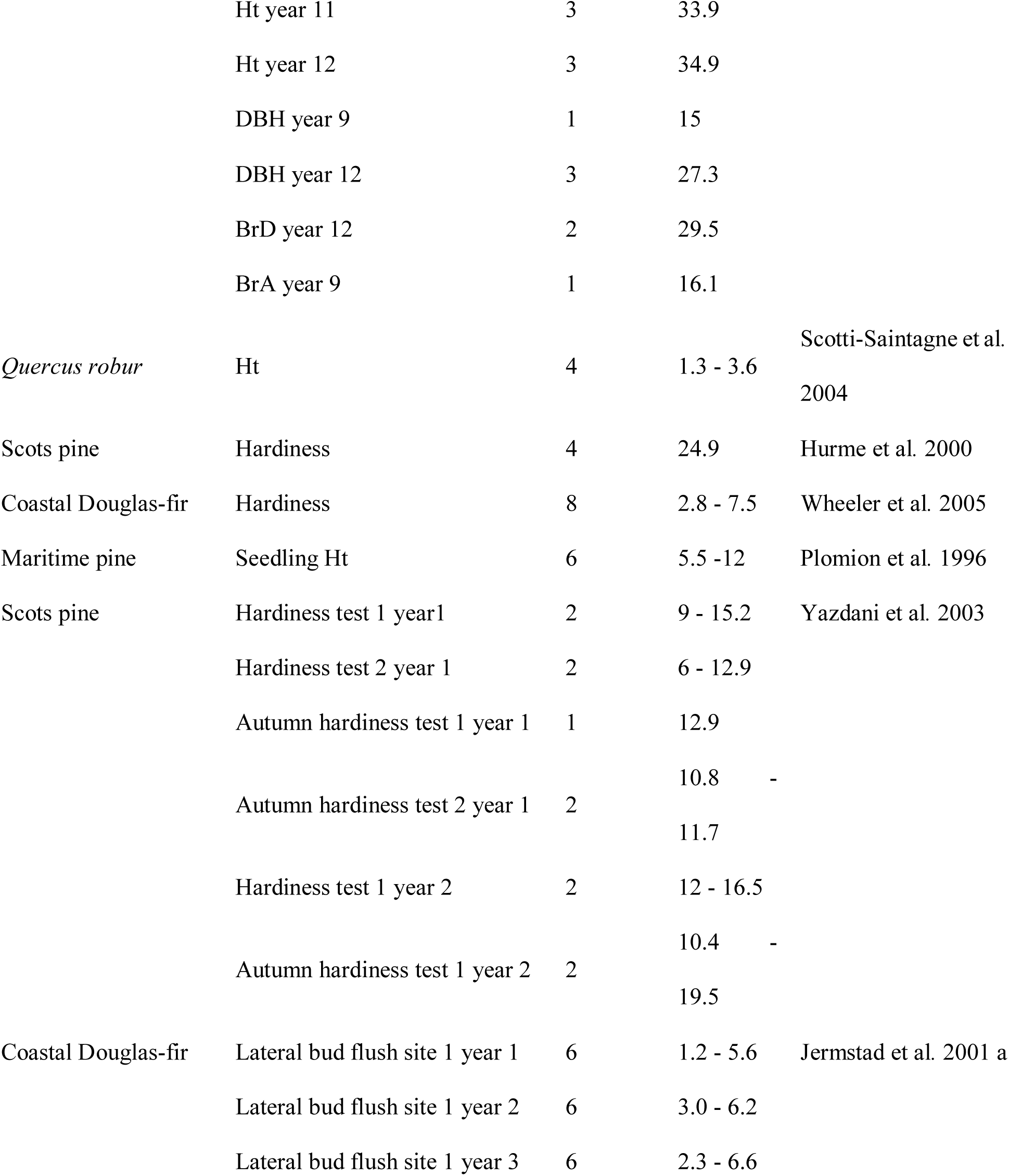
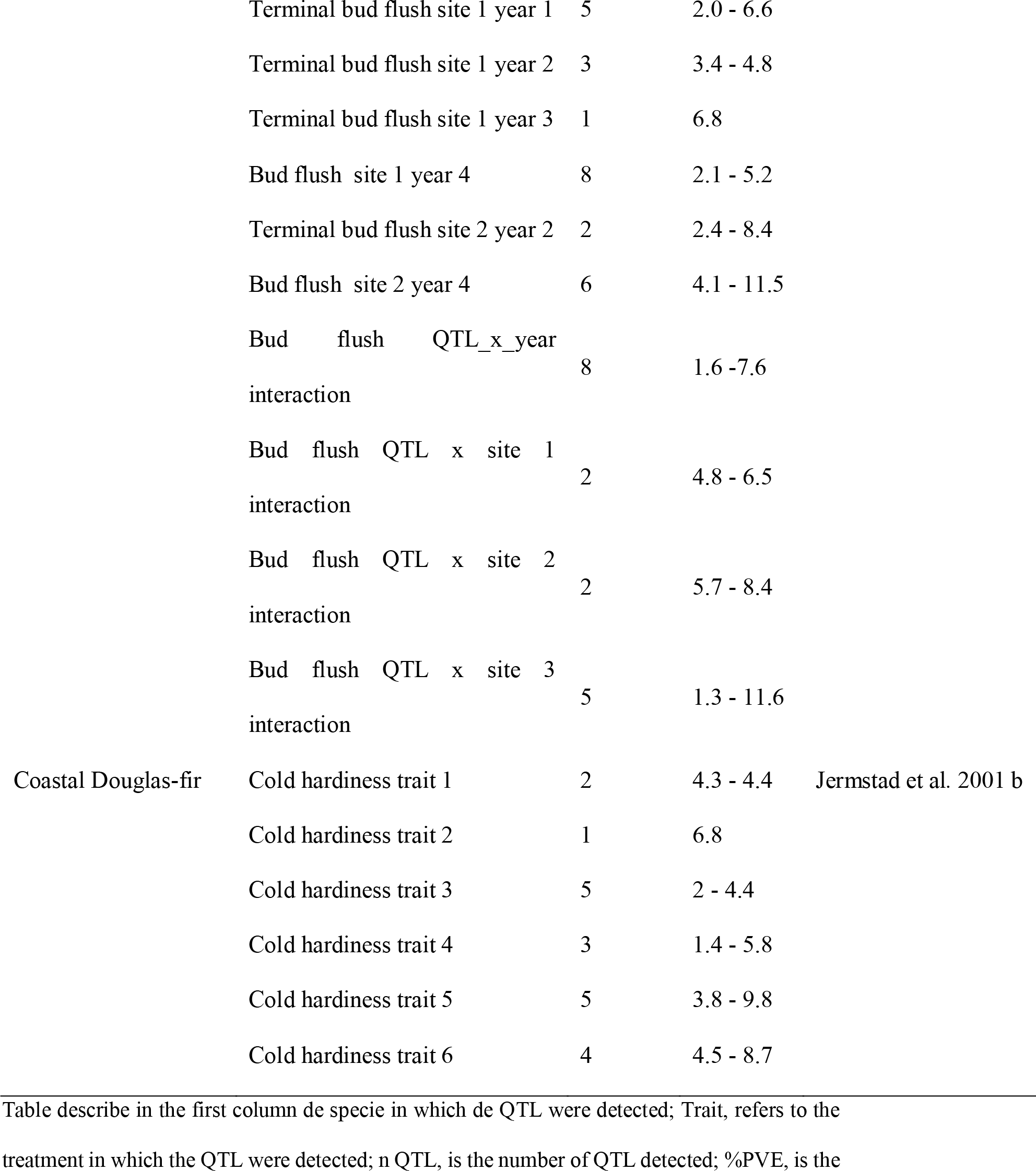
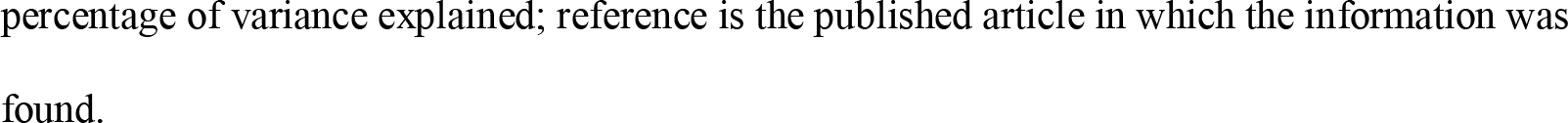
QTL detected in the literature for growth and adaptive traits

### Pleoiotropic effects

Pleiotropy is a complex phenomenon that refers to a single gene affecting multiple traits (He and Zhang, 2006). We have detected two SNP-QTL that may have pleiotropic effects. 0_7009_01-358 is significantly associated with FP, Ct and W and axs_47_502 is significantly associated with CO and Ct. Pleiotropic effects have previously been described in the conifer literature. For example, Pelgas et al. (2011), reported one possible common gene for budset, bud flush and height growth, and another for budset and height. Lerceteau et al. (2001) found one gene in common between DBH and BrD. Hurme et al. (2000) also found one gene with a possible pleiotropic effect between budset and frost hardiness. In the present work, it is however, not possible to discern between pleoiotropic effects and the action of two or more loci in the proximity. It has been described in animal breeding literature, that the possibitity of pleiotropic effects are hidding other effects such us linkage of separate loci (Johnsson et al. 2014; Wright et al. 2010). However, in our case we did not detect any significant correlation between the phenotypic traits for which we have detected pleiotropic effects.

### QTL age stability and QTLxE

We detected Ht and Ct QTL significantly associated to the same trait across several tree ages. This indicates that some of the QTL detected in this study could be considered strong candidates for early selection to certain traits based on their cross-age stability. High age-age correlations have been detected for DBH and Ht, in *Larix kaempferi* (Lai et al. 2014). In the literature, Ct has been shown to exhibit a moderate age-age correlation in some *Rhododendron* populations (Lim et al. 2014), whilst in *Phellodendron sachalinense* a high correlation of 0.91 was observed between 34 months old seedlings and 35 year old parent (McNamara and Pellett, 2000).

In our study, we could test the effect of environmental heterogeneity between-sites (i.e., different trials for the same families) and we could also compare the results of individual site versus MET analysis. The lack of reproducibility of the QTL across sites could be a consequence of environmental heterogeneity and G×E interactions observed among our study sites. Jansson (2007) and Persson et al. (2010) reported that the heritabilities of height and survival could vary significantly at different Scots pine progeny tests. Freeman et al. (2013) and Bartholomé et al. (2013) also found clear evidences of QTL×E interaction for growth traits in *Eucalyptus globulus* and *Eucalyptus* hybrids respectively. Rae et al. (2008) have shown that the environment influences the detection of QTL in three different locations in Europe, for bioenergy traits in *Populus* hybrids. In conifers, evidence of QTL×E interactions has been found for wood specific gravity in *Pinus taeda* L. (Groover et al. 1994).

### Significant SNP molecular functions

Several signifincant SNP-QTL may encode for known proteins. The best BLAST hit for 0_7009_01-358 is a kelch-motif containing protein. The kelch motif is an evolutionarily-widespread sequence motif of 44-56 amino acids in length involved in protein-protein interactions and the proteins that contain this motif are involved in the regulation of the circadian clock, in brassinosteroid modulation or in phenylpropanoid biosynthesis, among other functions (Zhang et al. 2013). The molecular and functional nature of such motif seems congruent with the pleiotropic nature of 0_7009_01-358. Another SNP-QTL involved in the control of CO and Ct is axs_47_502 that encodes for an UDP-apiose/xylose synthase. This enzyme catalyzes the NAD + -dependent conversion of UDPD-glucuronic acid to UDP-D-apiose and UDP-Dxylose (Grisebach and Schmid, 1972) and it is involved in the plant cell wall formation. UDP-Dxylose involvement in the pleiotropic control of CO and Ct may be the result of multiple genes in linkage disequilibrium within this single QTL. For example, fine QTL-mapping in rice identified multiple genes associated to the same QTL region (Lim et al. 2014). The 2_10212_01-241 encodes for a glutathione transferase located at the chloroplast. The involvement of this gene in stress regulation (Sappl et al. 2009) seems compatible with this SNP-QTL’ association with frost tolerance (Ct). The a3ip2_387 encodes for the ABI3-interacting protein 2 (ABI3), a transcription factor of the abscisic acid signal transduction pathway (Koornneef et al. 1984). In conifers, dormancy and frost tolerance are interconnected processes that share common molecular mechanisms with *Arabidopsis* (Welling and Palva, 2006). Considering the role of ABI3 in frost tolerance (Tamminen et al. 2001) and dormancy (Nambara et al. 1995) in *Arabidopsis*, it could be postulated that a possible involvement of ABI3 in frost tolerance also occurs in conifers, a hypothesis which is reinforced by the known association of this gene in seed development and dormancy in conifers (Zeng et al. 2013). The 2_10352_02-413 QTL’ best BLAST is a nucleotide sugar epimerase, which is involved in photosynthesis membrane biogenesis and its overexpression leads to growth acceleration in *Arabidopsis* (Li et al. 2011), thus supporting its role in conifer growth. The best BLAST hit result for QTL 2_6731_01-230 was a putative F-box protein GID2. F-box protein, gibberellin-insensitive dwarf2 (GID2), mediates the action of phytohormone gibberellin (GA) in the control of growth and development in plants (Gomi et al. 2004). This gives credibility to the SNP-QTL association with growth (Ht) found in this study. We also found a significant association between GI_f1_334 that encodes for gigantean (GI) with vitality (Vt). GI is a very well characterized gene involved in multiple physiological processes including stress responses to cold and drought in *Arabidopsis* (reviewed by Mishra and Panigrahi, 2015). This could be interpreted as a validation of GI involvement in vitality mediating processes in conifers. 0_11919_01-122, 2_9603_01-344, 0_17247_02-266 and CL2495Contig1_03-101 were not annotated at the time of this study.

## Conclusions

The identification of QTL associated with different growth, quality and adaptive traits through phenotypic measurements and EBVs, has resulted in the detection of QTL linked to 18 AFLPs and 12 SNPs. 62 significant QTL were detected for phenotype based traits, probably due to the larger number of phenotype based traits measured compared with the EBV-based traits. The levels of %PVE of the QTL ranged from 1.7 to 18.9. Multiple QTL were significantly associated to the same trait across ages, therefore those QTL would be considered as the most promising candidates for early selection. We detected environmental heterogeneity between sites and genotype-by-site interactions, which could be the main reason behind the absence of reproductitility of some QTL across sites. We detected Ht and Ct QTL stable across several tree ages, thus indicating that some of the QTL detected in this study could be considered strong candidates for early selection for certain traits based on their cross-age stability.

## Acknowledgements

We are thankful to Ross Whetten for guidance in the development and genotyping of AFLP markers, to Outi Savolainen for the SNP data, to Sonja Kujala for the information on SNP annotation, and to Skogforsk that planted and maintained the field material, and also for the help with filed measurements. We also grateful to Anders Fries and René Fernández for the field sampling, and, Dr. Fikret Isik, Dr. Zhi-Qiang Chen and Dr. Arthur Gilmour for their assistance in the statistical analysis. We are thankful to John Baison for his help with the English language review. This study was supported by the Research School in Forest Genetics and Breeding at the Swedish University of Agricultural Sciences, Knut och Alice Wallenbergs foundation, EVOLTREE EU project and PROCOGEN EU project.

## Data archiving

Phenotypic data and pedigree information are available in supplementary materials (S1). Genotype datasets are available in files S3 and S4 at Li et al. (2014). Sorting and mapping of the marker data are available at file S5 of Li et al. (2014). The authors state that all the other data necessary to support the conclusions of this work are fully represented within the article.

